# Regulatory interactions between Vax1, Pax6, and miR-7 regionalize the lateral Ventricular-Subventricular Zone during post-natal Olfactory Bulb neurogenesis in mice

**DOI:** 10.1101/2020.04.24.059428

**Authors:** Nathalie Coré, Andrea Erni, Pamela L. Mellon, Hanne M. Hoffmann, Christophe Béclin, Harold Cremer

**Affiliations:** Aix Marseille University, CNRS, IBDM, Campus de Luminy, Marseille, France; Department of Obstetrics, Gynecology, and Reproductive Sciences and the Center for Reproductive Science and Medicine, University of California, San Diego, La Jolla, California 92093-0674, USA

## Abstract

Several subtypes of interneurons destined for the olfactory bulb are continuously generated after birth by neural stem cells located in the ventricular-subventricular zones of the lateral ventricles. Future neuronal identity depends on the positioning of pre-determined neural stem cells along the ventricle walls, which, in turn, depends on delimited expression domains of transcription factors and their cross regulatory interactions. However, mechanisms underlying positional identity of neural stem cells are still poorly understood. Here we show that the transcription factor Vax1 controls the production of two specific neuronal sub-types. First, it is directly necessary to generate Calbindin expressing interneurons from ventro-lateral progenitors. Second, it represses the generation of dopaminergic neurons by dorso-lateral progenitors through inhibiting Pax6 expression in the dorso-lateral wall. We provide evidence that this repression occurs via activation of microRNA miR-7, targeting Pax6 mRNA.

## Introduction

In the postnatal and adult rodent forebrain, new interneuron precursors are continuously generated by neural stem cell populations along the walls of the lateral ventricles. After their amplification in the subventricular zone (SVZ) and long-distance migration via the rostral migratory stream (RMS), they are added to the preexisting circuitry of the olfactory bulb (OB) (Alvarez-Buylla and Garcia-Verdugo, 2002; Whitman and Greer, 2009; Platel et al., 2019).

New OB interneurons produced in this SVZ-RMS-OB neurogenic system show a wide spectrum of phenotypic diversity at the levels of morphology, terminal position, connectivity, and neurotransmitter use (Whitman and Greer, 2007). Lineage studies demonstrated that this diversity relies on the existence of defined microdomains of predetermined neural stem cells in the ventricular-subventricular zone [V-SVZ (Merkle et al., 2007; Ventura and Goldman, 2007; Lledo et al., 2008; Fiorelli et al., 2015; Chaker et al., 2016)].

A key question concerns the molecular mechanisms underlying this diversity. It has been shown that Gli1 activation by Sonic Hedgehog (SHH) is necessary to generate Calbindin-expressing periglomerular neurons (CB-N) in the ventral aspect (Ihrie et al., 2011). Moreover, the zinc-finger transcription factors (TF) Zic1 and Zic2 act as inducers of Calretinin (CR)-expressing GABAergic interneurons in the dorso-septal region (Tiveron et al., 2017). However, the high diversity of the postnatal generated interneuron subtypes suggests that more complex molecular cascades and cross regulatory interactions are put in place to define the stem cell compartment at the necessary resolution.

This is exemplified by the regulation of a group of OB interneurons that, in addition to GABA, use dopamine as their main neurotransmitter (DA-N). This neuron type is generated by neural stem cells located in the dorsal and dorso-lateral aspects of the ventricular wall. Pax6 is a key determinant of this cell type (Hack et al., 2005; Kohwi et al., 2005) and is expressed in the entire lineage from stem cells to neurons (Hack et al., 2005; de Chevigny et al., 2012b). In addition to this positive transcriptional regulation, posttranscriptional mechanisms have been shown to be crucial for the negatively control of DA-N production. Indeed, Pax6 mRNA expression in the postnatal ventricular wall is not restricted to the DA-N producing dorsal progenitor pool, but extends far into the lateral region where other cell types, including purely GABAergic granule cells and CB-N, are produced. However, the presence of the microRNA miR-7 in a Pax6-opposing ventro-dorsal gradient precludes Pax6 mRNA translation and restricts protein expression, and consequently DA-N phenotype, to the dorsal region (de Chevigny et al., 2012a). Thus, complex molecular events implicating transcriptional and post-transcriptional control mechanisms underlie the functional diversity of OB interneurons.

The Ventral Homeodomain Protein 1 (Vax1), an intracellular mediator of SHH signaling, is expressed in ventral territories of the developing mouse forebrain as well as in ventral aspects of the developing eye (Hallonet et al., 1999; Ohsaki et al., 1999). In both systems, the expression pattern of Vax1 is complementary to that of Pax6, and genetic studies provided evidence for cross regulatory interactions between both factors (Hallonet et al., 1998; Bertuzzi et al., 1999; Hallonet et al., 1999; Stoykova et al., 2000; Baumer et al., 2002; Mui et al., 2005).

At embryonic stages, constitutive Vax1 mutants show a strong decrease in GABAergic interneurons in the developing neocortex, indicating an essential function in their generation (Taglialatela et al., 2004). Vax-1 homozygous mutants die generally at perinatal stages and only few “escapers” survive for a few weeks after birth. In these animals, the entire postnatal SVZ-RMS-OB neurogenic system is severely compromised, showing accumulation of precursors in the SVZ and severe disorganization of the RMS (Soria et al., 2004), altogether precluding a detailed analysis of Vax1-function at later stages.

In an attempt to understand the regulatory cascades underlying postnatal OB interneuron diversity, we identified Vax1 as a potential candidate. Indeed, based on a high-resolution gene expression screen, comparing the postnatal pallial and subpallial OB lineages, we found that Vax1 mRNA is present in a ventro-dorsal gradient along the lateral wall of the forebrain ventricles. Functional studies using conditional mutants demonstrate that Vax1 is essential for the generation of CB-N by the ventral stem cell pool. Moreover, we show that Vax1 acts as negative regulator of DA-N OB fate via downregulation of the pro-dopaminergic factor Pax6. Finally, we provide evidence that this repressor function is, at least in part, mediated by induction of miR-7.

## Results

### Vax1 is expressed in a ventro-dorsal gradient along the lateral ventricle

We investigated gene expression during postnatal OB neurogenesis by in vivo electroporation of neural stem cells in the lateral and dorsal aspects of the forebrain lateral ventricle at postnatal day 1 (P1), followed by the isolation of homotypic cohorts at different time points by microdissection and FACS. Microarray analyses provided detailed insight into gene expression changes between the two neurogenic lineages (“in space”) and during the progression from stem cells to young neurons (“in time”; Figure 1A; for detail see Tiveron et al, 2017).

**Figure 1.**
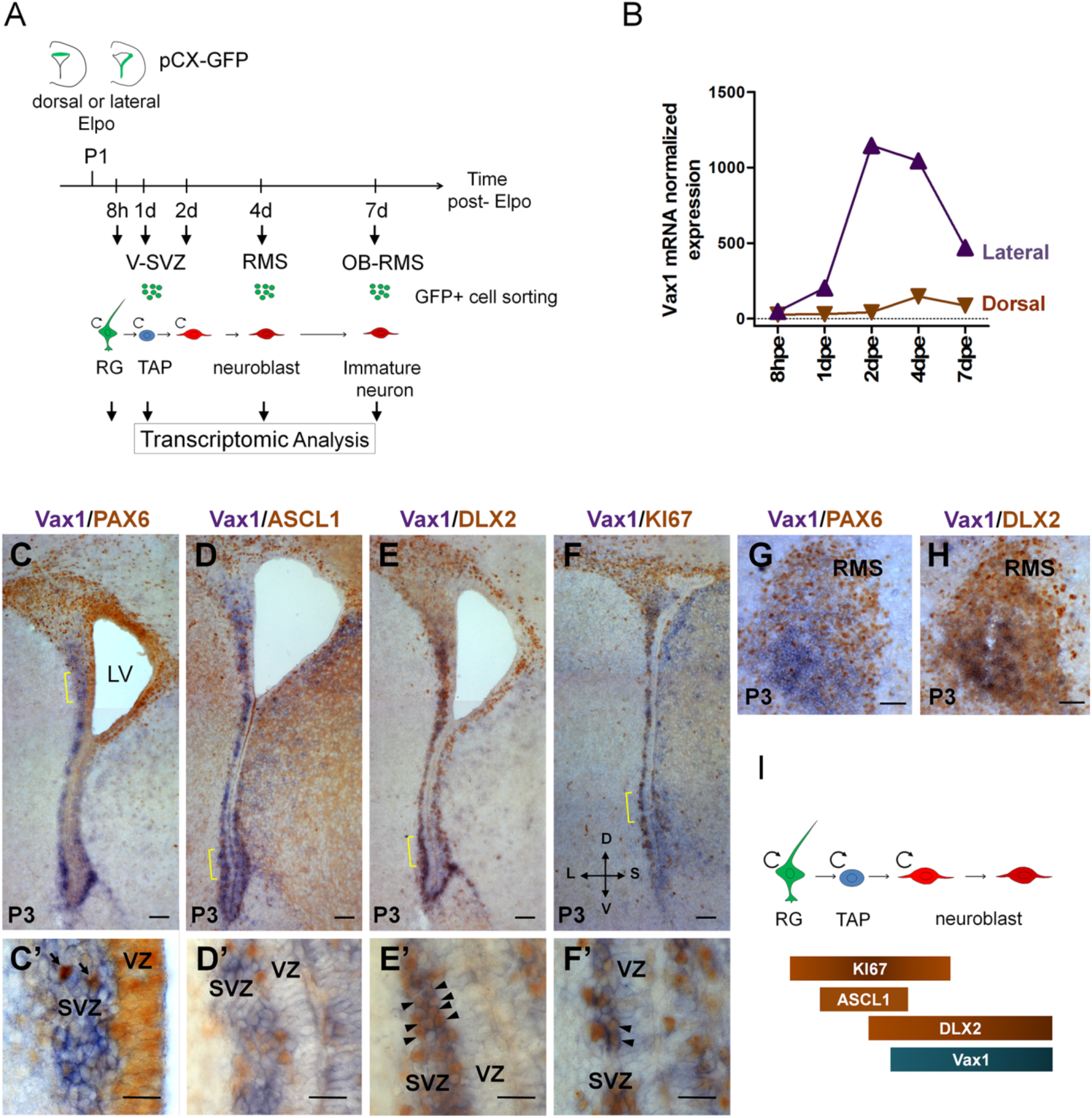
Vax1 is expressed in the lateral V-SVZ. (A) Representation of the strategy used for transcriptomic analysis in time and space in the dorsal and lateral lineages. pCX-GFP plasmid was introduced into neural stem cells (NSCs) residing within the dorsal or lateral V-SVZ and GFP-positive cells were isolated by FACS at different time points after electroporation (Elpo). The mRNA content was analyzed by µarray (Tiveron et al., 2017). (B) Quantification of Vax1 mRNA expression detected by µarray analysis in dorsal (brown) and lateral (purple) progenies during neurogenesis. (C-H) In situ hybridization revealing Vax1 mRNA (in blue) combined with immuno-histochemistry using antibodies detecting (in brown) PAX6 (C, C’, G), ASCL1 (D, D’), DLX2 (E, E’, H) or KI67 (F, F’) proteins in the V-SVZ (C-F) or RMS (G, H) at post-natal day 3 (P3). (C’-F’) High magnification of cellular staining in the V-SVZ (area indicated by the yellow bracket in C-F). Arrows (C’): examples of strong PAX6 only positive cell in the dorso-lateral SVZ; blue staining underneath labels cells from distinct plane. Arrow heads (E’, F’): double positive cells for DLX2 and KI67 respectively. (I) Schematic representation of gene expression profile in sequential cell types of the neurogenic sequence. Circular arrow indicates proliferating cells. LV: lateral ventricle, RG: radial glia, TAP: transit amplified precursor, VZ: ventricular zone, SVZ: sub-ventricular zone. D: dorsal, L: lateral, S: septal, V: ventral. Scale bars: 100µm (D-G), 20µm (D’-G’), 50µm (I-H).

These analyses showed that Vax1 was confined to the neurogenic lineage derived from the lateral ventricular wall (Fig. 1B). Vax1 mRNA was induced at low levels at 1 day post-electroporation (dpe) when most GFP positive cells were transit amplifying precursors [TAPs, (Boutin et al., 2008); Figure 1B]. Expression strongly increased at 2dpe and remained stably high at 4dpe, when most cells were migratory neuronal precursors, before steeply decreasing at 7dpe when cells arrived in the OB and emigrated from the RMS to invade the granule cell (GCL) and the glomerular (GL) layers (Tiveron et al., 2017). In comparison, isolates from dorsally electroporated brains showed no obvious Vax1 mRNA expression over all analyzed time points (Figure 1B).

Next, we investigated Vax1 co-expression with known markers of defined neuronal subsets or differentiation stages in the postnatal V-SVZ. In the absence of a Vax1 antibody that provided reliable signals on post-natal tissue sections, we combined in situ hybridization for Vax1 mRNA with immunohistochemistry for the proteins PAX6, ASCL1, KI67, or DLX2 (Figure 1C-H). Vax1 mRNA always showed a gradient-like distribution along the lateral ventricular wall, with highest expression in the ventral-most aspect and extending far into dorsal regions (Figure 1C,D,E,F). Fainter expression was observed along the septal wall. In the lateral wall, Vax1 mRNA was excluded from the VZ but expressed in the underlying SVZ (Figure 1C’,D’,E’,F’). Vax1 mRNA in the lateral wall was generally non-overlapping with PAX6, a marker for the dorsal stem cell pool and a determinant for the DA-N lineage (Figure C,C’). Vax1 mRNA positive cells rarely expressed ASCL1, a marker for TAPs (Figure 1D,D’), but were labeled with a DLX2 antibody (Figure 1E,E’; arrowheads), confirming the preferential expression of Vax1 in migratory neuronal precursors (Doetsch et al., 2002) (Figure 1B,I). Finally, about one third of the Vax1+ cells in the SVZ expressed the proliferation marker KI67 (Figure 1F,F’, arrowheads), which likely correspond to mitotic neuroblasts. In the RMS, Vax1 mRNA was confined to the ventral part, complementary to PAX6 immunostaining (Figure 1G) and co-localized with DLX2 (Figure 1H).

Altogether, the combination of microarray studies in defined neuronal lineages and histological approaches leads to the conclusion that Vax1 is expressed in the lateral and ventral SVZ in a subset of proliferating precursors and in most migratory neuroblasts, the latter maintaining expression during their migration in the RMS.

### Vax1 is necessary for the generation of Calbindin interneurons

Previous work demonstrated that Calbindin-positive GABAergic interneurons (CB-N) destined for the GL are generated from the ventral-most region of the anterior lateral ventricles (LV), and SHH signaling has been implicated in their specification (Merkle et al., 2007; Ihrie et al., 2011). As Vax1 is strongly expressed in this area, and has been shown to act as an intracellular mediator of SHH signaling (Take-uchi et al., 2003; Furimsky and Wallace, 2006; Zhao et al., 2010), we first asked if this TF is implicated in the generation of CB neurons.

Vax1 conditionally mutant mice (Vax1cKO) (Hoffmann et al., 2016) were bred to R26tdTomato mice to monitor CRE-induced recombination and to follow the distribution and fate of mutant and control cells over time. We used postnatal in vivo brain electroporation to express CRE protein in the lateral wall (Figure 2A). Since targeting of the Vax1-positive ventral region of the ventricular wall with DNA-based expression constructs is inefficient, we used CRE mRNA, that is highly efficient for the transfection of stem cells along the entire wall, including the most ventral aspect (Bugeon et al., 2017). Animals were electroporated at P0 and analyzed 15 days later, when labeled neurons reached the OB and integrated into the GCL and GL (Figure 2B). Quantification of labelled neurons in the GCL revealed no differences between control and mutant mice (Figure 2C), demonstrating that generation of the granule cells was not dependent on Vax1 function. However, there was a significant loss of CB-positive neurons in the GL of the OB at 15dpe (Figure 2D,E).

**Figure 2.**
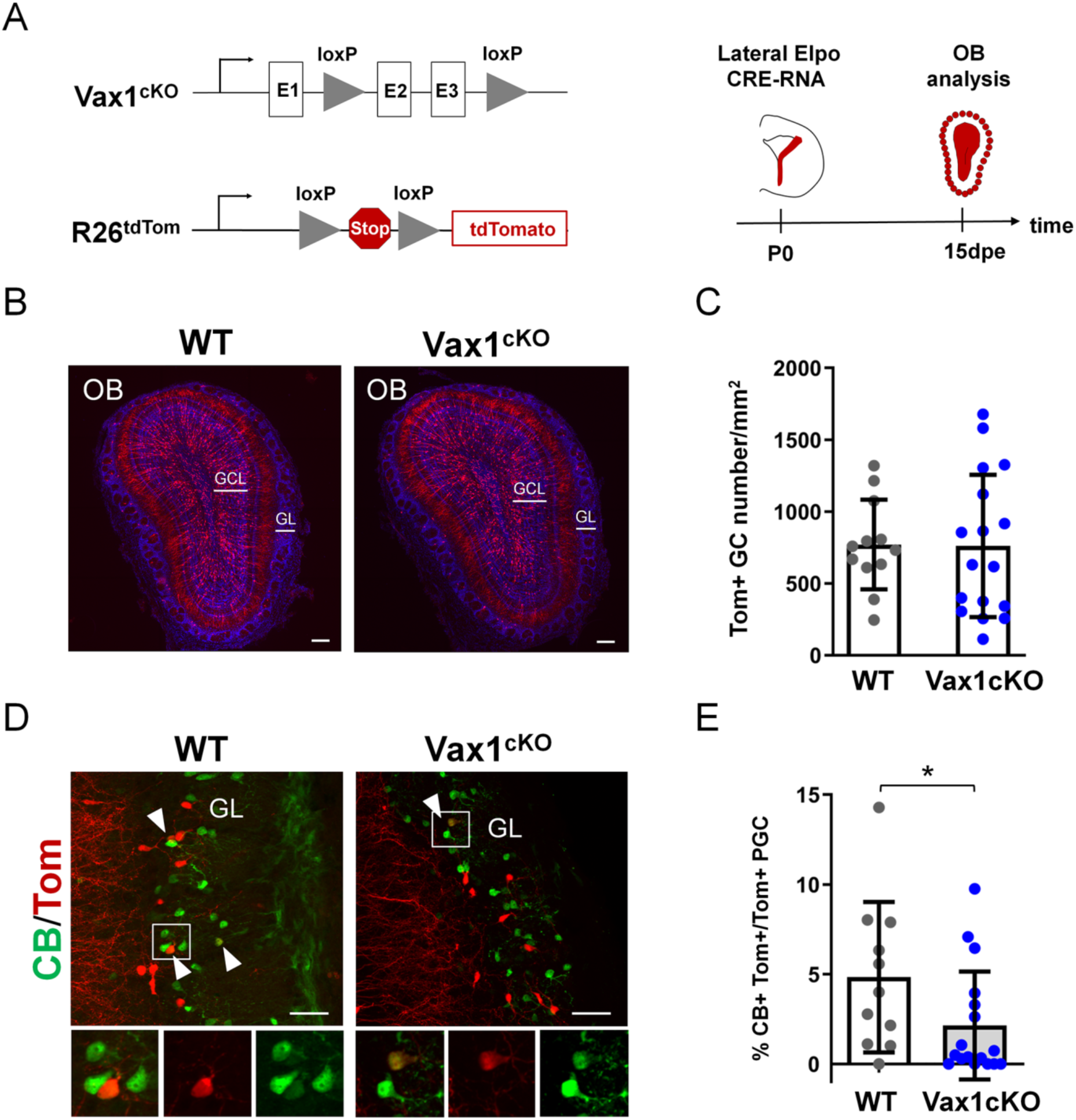
Vax1 is necessary for the production of Calbindin-positive interneurons in the olfactory bulb. (A) Representation of the Vax1 conditional allele (Vax1cKO) combined with the inducible reporter tdTomato allele at the Rosa26 locus (R26tdTom). Right panel: strategy used to recombine the Vax1 mutant allele in the V-SVZ cells in the lateral wall at post-natal day 0 (P0). TdTomato (Tom)-positive cells were analyzed 15 days post electroporation (dpe) in the olfactory bulb (OB). (B) Images showing the distribution of Tom+ cells (in red) in the OB at 15dpe in control and mutant brains. Nuclei (in blue) are stained by Hoechst. (C) Quantification of granule cells (GC) number in the OB GCL in both conditions. Data are shown as means ± SD, dots represent individual animals. WT: n= 12, Vax1cKO: n= 17. (D) Images showing Calbindin+ (in green) and Tom+ cells in the GL at 15dpe in control (n=11) and Vax1 mutant (n=17) brains. Arrow heads indicate double stained neurons. High magnification of representative of double positive cells is shown below. GCL: granule cell layer, GL: glomerular layer. * p ≤ 0.05. Scale bars: 200µm (B), 50µm (D).

Thus, Vax1 expression in the ventro-lateral-derived neurogenic lineage is necessary for the correct generation of CB-N in the GL.

### Vax1 regulates Pax6 during OB neurogenesis

In situ hybridization indicated that Vax1 mRNA was expressed in a ventro-dorsal gradient (Figure 1). To confirm the existence of such a gradient, we micro-dissected V-SVZ tissue from the dorsal, dorso-lateral, and ventro-lateral regions of the ventricular walls and subjected the isolates to RT-qPCR analyses for Vax1 mRNA. In agreement with the histological data, Vax1 mRNA showed a steep ventro-dorsal gradient, opposed to, and partially overlapping with, the well-described localization of Pax6 mRNA, that extends dorso-ventrally (Figure 3A) (de Chevigny et al., 2012a). This observation appeared significant for two main reasons. First, several studies provided evidence that Vax1 can negatively regulate Pax6 expression during development (Bertuzzi et al., 1999; Hallonet et al., 1999; Mui et al., 2005). Second, repression of Pax6 translation along the lateral wall is necessary to confine Pax6 protein, and consequently dopaminergic OB neuron phenotype, to the dorsal stem cell pool (de Chevigny et al., 2012a).

**Figure 3.**
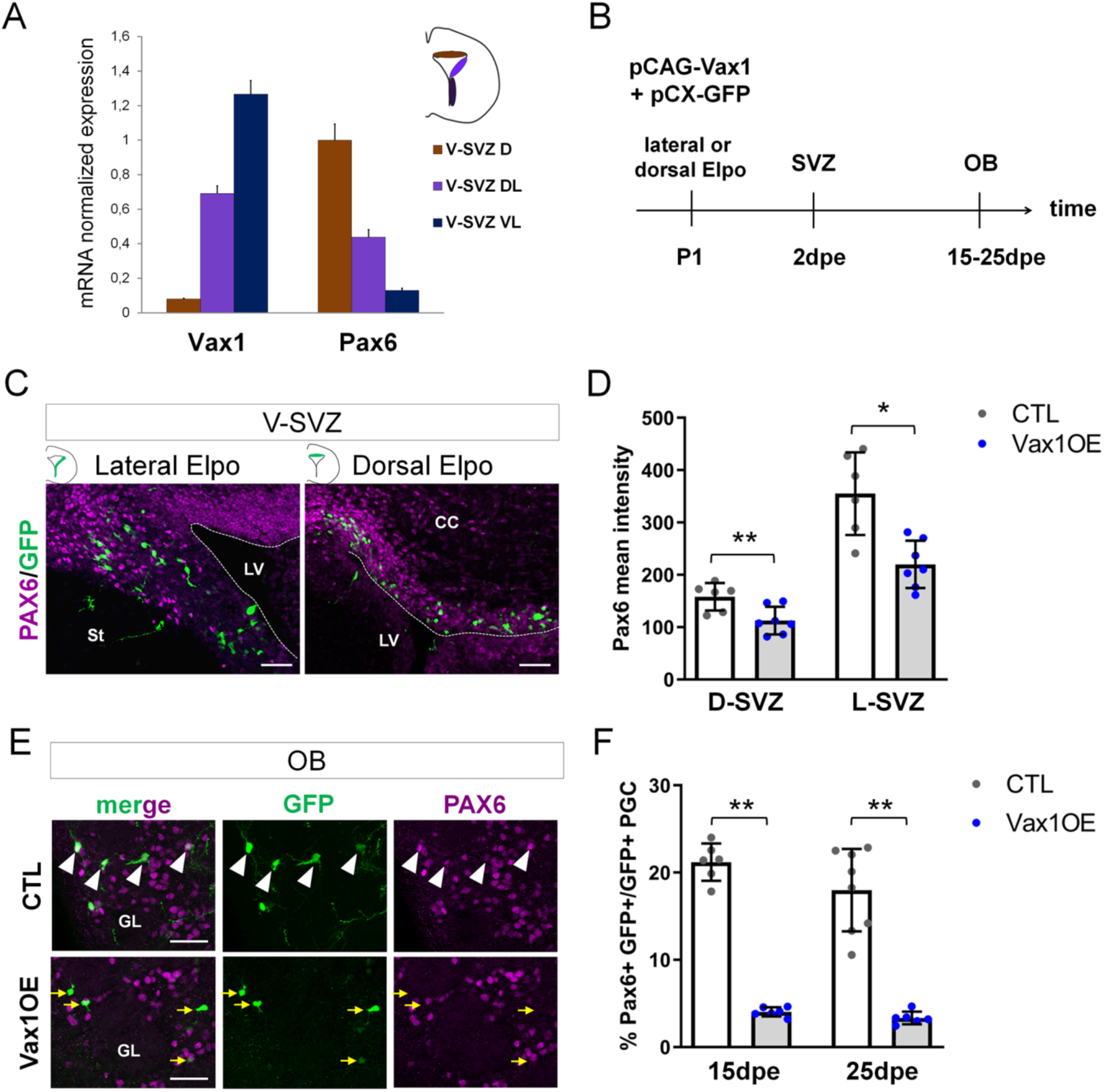
Vax1 inhibits PAX6 expression in the V-SVZ and the OB. (A) Quantitative RT-PCR revealing Vax1 and Pax6 gene expression in cells dissected out from 3 distinct areas of the V-SVZ. D: dorsal, DL: dorso-lateral, VL: ventro-lateral. (B) Strategy design for the Vax1 gain-of-function experiment. The Vax1 expressing plasmid (pCAG-Vax1) was introduced into lateral or dorsal progenitors in combination with pCX-GFP by electroporation at P1. Brains were analyzed at different time points in the V-SVZ or the OB. (C) Representative images showing simultaneous expression of PAX6 and GFP proteins in dorsal or lateral lineage in the V-SVZ. (D) Quantification of PAX6 mean intensity in control or Vax1-overexpressing (OE) V-SVZ GFP+ cells from dorsal (D, n=6 for the control, n=7 for Vax1 condition) or lateral (L, n=6 for the CTL, n= 7 for Vax1 condition) walls. (E) Images showing simultaneous expression of PAX6 and GFP in the OB glomerular layer of control or Vax1OE brains. Arrow head: double GFP/PAX6-positive cells; yellow arrow: GFP only cells. (F) Quantification of GFP+PAX6+ neurons in the OB GL at 15dpe (n=6 for the CTL, n=6 for Vax1OE) and 25dpe (n=8 for the CTL, n=6 for Vax1OE). PGC: periglomerular cell. * p ≤ 0.05, **p ≤ 0.01. All scale bars: 50µm.

Based on this information we hypothesized that Vax1 is implicated in Pax6 down-regulation in the postnatal SVZ. To test this, we overexpressed Vax1 in the dorsal and lateral neurogenic lineages by electroporation and investigated the impact on Pax6 expression. A Vax1 expression plasmid (pCAG-Vax1), or an empty control vector, was co-electroporated with pCX-GFP into either the dorsal or the lateral ventricular wall (Figure 3B). Two days later, animals were sacrificed and Pax6 immunostaining intensity in GFP positive cells was measured in the dorsal and dorso-lateral SVZ (Figure 3C,D). GFP-positive cells generated in both compartments showed a significant decrease in Pax6 expression levels at this early time point (Figure 3D).

Then we asked if Pax6 expression was affected at late time points, after the arrival of newborn neurons in the OB GL. Indeed, 15 and 25 days after pCAG-Vax1 electroporation into the dorsal wall, the proportion of GFP-positive neurons that showed Pax6 expression was reduced by over 80% (Figure 3E,F).

We conclude that Vax1 has the capacity to act as a negative regulator of Pax6 expression during postnatal OB neurogenesis.

### Vax1 negatively regulates dopaminergic phenotype

Dopaminergic neurons in the OB GL are derived from the dorsal and latero-dorsal aspects of the ventricle walls (Merkle et al., 2007; Fernandez et al., 2011; de Chevigny et al., 2012a) and Pax6 expression is necessary and sufficient for the acquisition of this neurotransmitter phenotype (Hack et al., 2005; Kohwi et al., 2005). We asked if Vax1 over-expression in Pax6-positive cells was sufficient to inhibit the generation of dopaminergic neurons in the OB.

We first targeted the dorsal compartment (Figure 4A-D), where the majority of dopaminergic neurons are generated (Fernandez et al., 2011; de Chevigny et al., 2012a). Ectopic expression of Vax1 led to a significant loss of TH/GFP-positive neurons 15 days later in the OB (Figure 4B). Loss of TH-positive cells was robust over time and could be observed at 25 and 60 dpe (Figure 4C). Two observations pointed towards the specific loss of dopaminergic neurons. First, the number of the second major identified neuron type that is generated in the dorsal ventricular wall, CR-N of the glomerular layer (Fernandez et al., 2011; Tiveron et al., 2017), was unaffected by Vax1 expression (Figure 4B), arguing against a fate shift towards this neuron type (Tiveron et al., 2017). Second, the density of total GFP+ cells in the GL of Vax1-electroporated animals was significantly reduced (Figure 4D) whereas the number of GFP+ granule cells was not affected (Figure 4-supplement figure 1A). As dopaminergic neurons represent a substantial population of all dorsal generated PG neurons such an overall loss is coherent with a loss of the DA-N subtype.

**Figure 4.**
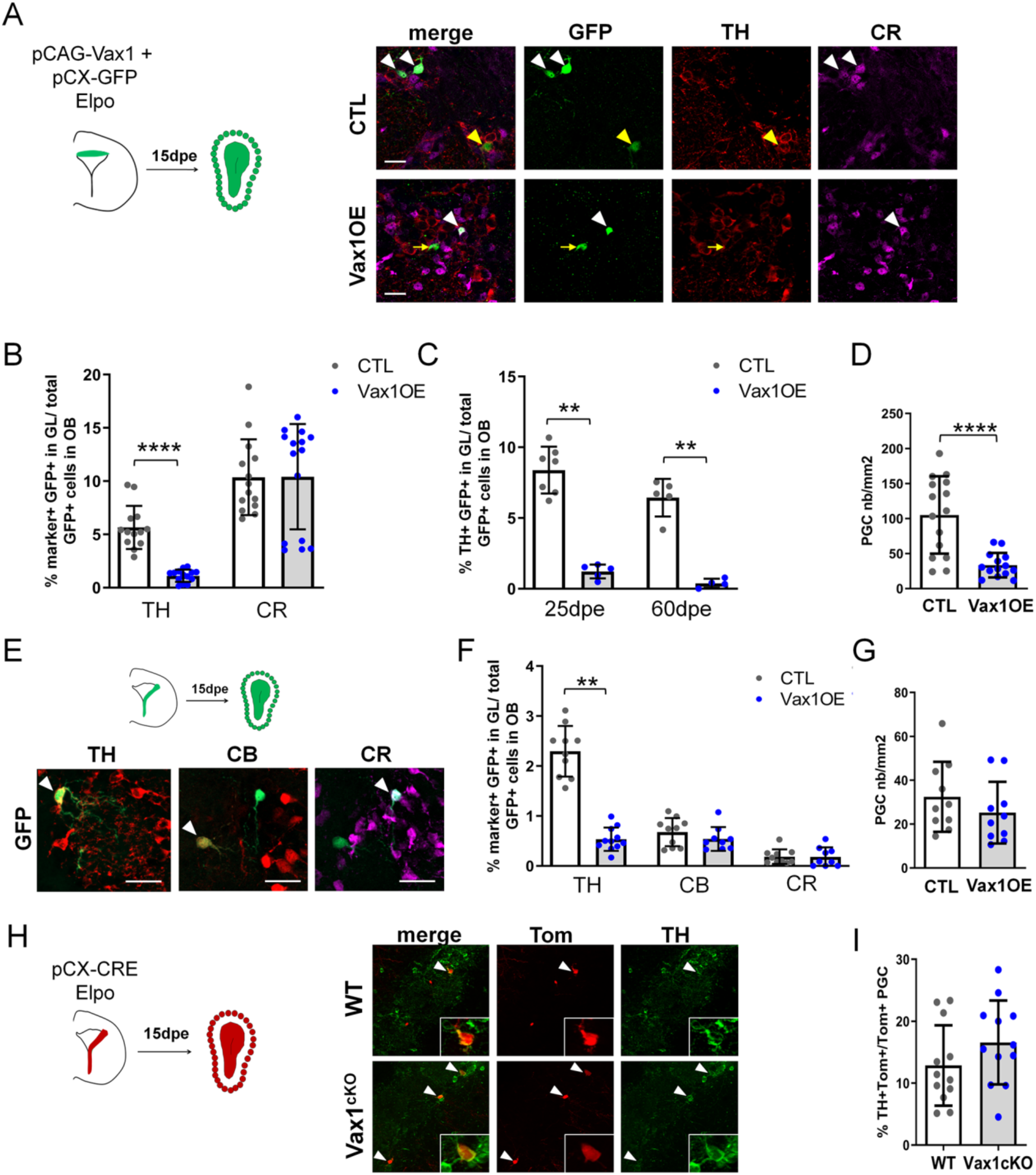
Over-expression of Vax1 in V-SVZ neural stem cells inhibits dopaminergic phenotype. Experimental design (left) for the electroporation of NSCs in the dorsal wall with pGAC-Vax1 + pCX-GFP. Images (right) showing expression of tyrosine hydroxylase (TH) and Calretinin (CR) in the OB glomerular layer 15 days after electroporation in Vax1 over-expression (OE) and control brains. White arrow head: GFP/CR-double positive neuron, yellow arrow head: GFP/TH double positive neuron, yellow arrow: GFP-only cell. (B) Histogram showing the reduction of the density of GFP+ periglomerular cells (PGC) in the Vax1OE OB (CTL n=15, Vax1 n=14). (C) The quantification of TH+ and CR+/GFP-positive cells shows a large decrease of the proportion of dopaminergic neurons among the total GFP+ cells in the OB of Vax1 condition (TH n=14, CR n=15) compared to control (n= 13/14). (D) The reduction of the TH+ population is sustained with time as it is still observed at 25- (CTL n=7, Vax1 n=5) and 60- (CTL n=5, Vax1 n=4) days post electroporation. (E) Experimental design (left) for the electroporation of NSCs in the lateral wall with pGAC-Vax1 + pCX-GFP. Representative images (right) of immunostaining with TH, Calbindin (CB), and CR antibodies in the OB GL. Arrow head: example of double positive staining with GFP for each marker. (F) Histogram presenting the quantification of the three different neuronal populations among the GFP+ neurons in the OB of control (n=10 for each marker) or Vax1OE (TH n=11, CB and CR n=9) conditions. (G) Histogram showing the density of GFP + PGC in both conditions (CTL n=10, Vax1 n=10). (H) Lateral NSCs of Vax1cKO: rosa26tdTom brains were electroporated at birth with pCX-CRE and neuronal phenotype was analyzed in OB at 15 dpe. Representative images of TH staining in the GL of control or Vax1 deficient OB. Arrow head: GFP+ cells co-labelled with TH. Insert: high magnification of a double positive neuron. (I) Histograms presenting the percentage of TH+ neurons among Tom+ PGC (CTL: n=12, 3 independent litters; Vax1: n=12, 3 independent litters). A slight increase of the TH+ population was observed in absence of Vax1 compared to control but statistical test (Mann Whitney U test) failed to give significant p values (p= 0.16). **p ≤ 0.01, ****p ≤ 0.0001. All scale bars: 20µm except in H (50µm).

Next, we targeted the lateral ventricular wall (Figure 4E-G), where smaller but still significant numbers of DA-N are produced from a dorso-lateral stem cells pool (Figure 4F). Overexpression of Vax1 in the lateral wall induced a significant loss of TH-positive neurons in the GL 15 days later (Figure 4F). At the same time point, numbers of CB-N and CR-N were unchanged (Figure 4F), indicating again that no phenotypic switch towards these subtypes occurred. The density of GFP+ cells was also unaffected in both GL (Figure 4G) and GCL (Figure 4-supplement figure 1B). Thus, Vax1 overexpression specifically inhibits DA-N phenotype of newborn neurons in the OB.

We also investigated whether Vax1 loss-of-function had a positive impact on DA-N phenotype in the OB. NSCs along the lateral wall of Vax1cKO animals were electroporated with pCX-CRE (Figure 4H) and the proportion of TH-positive neurons in the OB was analyzed 15 days later. These analyses failed to show a significant increase in DA-N at a confidence level of p ≤ 0.05. However, we stably observed a tendency for an increase over independent electroporation experiments (3 experiments for each condition with a total of 12 animals/ condition, Figure 4I).

Together, these results show that Vax1 acts as a specific repressor of the DA-N phenotype. We conclude that Vax1, likely via regulation of Pax6, has the capacity to negatively control the generation of dopaminergic neurons for the OB. Moreover, these data also show that while Vax1 is necessary for the generation of CB-N, it is not sufficient.

### Vax1 induces miR-7 expression in the lateral wall

Previous work demonstrated that microRNA miR-7 is expressed in a ventro-dorsal gradient along the lateral ventricular wall and post-transcriptionally inhibits Pax6 protein expression. This interaction confines the generation of dopaminergic neurons to the dorsal aspect (de Chevigny et al., 2012a). As Vax1 and miR-7 are expressed in a similar gradient, we hypothesized that the repression of Pax6 protein expression by Vax1 is mediated by miR-7. To address this idea, we overexpressed Vax1 together with GFP in the lateral stem cell compartment and isolated GFP-positive cells two days later by microdissection, dissociation and flow cytometry cell sorting (Figure 5A). qRT-PCR analyses demonstrated that augmented Vax1 expression (Figure 5B) led to a strong increase in miR-7 levels (Figure 5C). Altogether, these results provide evidence that the negative impact of Vax1 on Pax6 expression is, at least in part, mediated via the positive regulation of miR-7.

**Figure 5.**
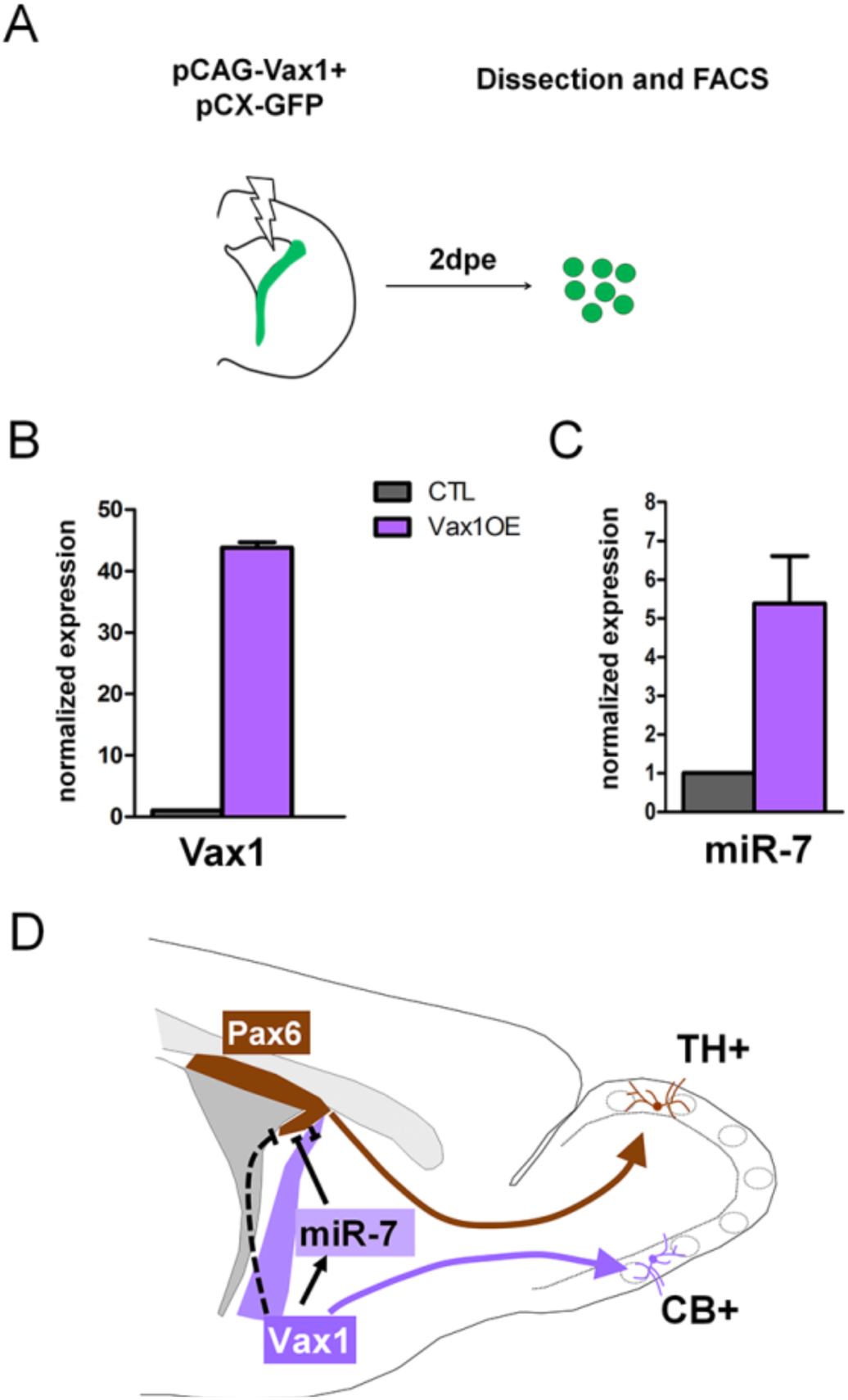
Vax1 induces the expression of miR-7 in the lateral V-SVZ. (A) Strategy used to determine the expression of microRNAs in Vax1-overexpressing progenitors. PCAG-Vax1 and pCX-GFP were simultaneously introduced into NSCs by electroporation of the lateral wall of post-natal P1 brains. Lateral V-SVZ was dissected out 2 days after electroporation and GFP+ cells were isolated by flow cytometry (FACS) to perform quantitative RT-PCR analysis. (B) Quantification of Vax1 mRNA level by qRT-PCR in control and Vax1OE conditions, normalized to beta-actin and reported in Vax1 condition as relative level to control, validating the overexpression of Vax1 after electroporation. (C) Quantification of miR-7 expression in both conditions. Expression level of miR-7 was normalized by invariant expression of microRNA let-7a and reported in Vax1 condition as relative level to control. Experiments in B and C were performed in triplicate and data were obtained from 2 technical replicates. (D) Model of cross-regulatory interaction between Vax1, miR-7, and Pax6 in the lateral V-SVZ to control the number of dopaminergic neurons generated by the neural stem cells regionalized in this aspect. This model is supported by our present data and previous work (de Chevigny et al., 2012a) where it was shown that miR-7 was required to inhibit PAX6 expression in lateral NSCs to produce the correct number of dopaminergic neurons in the post-natal OB. Here we propose that Vax1 acts upstream of miR-7 by positively regulating its expression and consequently inhibiting PAX6. However, it is possible that Vax1 also directly represses the expression of Pax6 mRNA (dashed line) by acting on its promoter (Mui et al., 2005). Additionally, Vax1 is required to generate Calbindin neurons from the ventral aspect of the lateral V-SVZ.

## Discussion

Here we show that Vax1 is strongly expressed in the ventral compartment of the lateral ventricle where it is necessary for the generation of CB-N. In addition to this actively phenotype-determining function, our data show that Vax1 acts as a negative regulator of Pax6, likely via the induction of miR-7, thereby restricting the generation of DA-N that are generated in a neighboring progenitor domain (Figure 5D) (de Chevigny et al., 2012a).

During nervous system development, determination of neuronal phenotype is controlled by the combinatorial expression of transcription factors (Flames et al., 2007; Guillemot, 2007; Lai et al., 2016). Moreover, cross regulatory interactions between such TFs have been shown to tightly define progenitor domains that generate specific neuron subtypes (Jessell, 2000; Sagner and Briscoe, 2019). Such spatial information has to be maintained during postnatal and adult stages and neuronal output has to be adapted to the needs of the ongoing neurogenesis in the OB. In agreement, regionalization of the postnatal stem cell compartment has been shown to depend on spatially restricted and combinatorial expression of TFs like, for example, Pax6, Emx1, Gsx1/2, Gli1/2, or Zic1/2 (Alvarez-Buylla et al., 2008; Weinandy et al., 2011; Fiorelli et al., 2015; Angelova et al., 2018).

CB-N are produced by NSCs positioned in the ventral aspect of the V-SVZ (Merkle et al., 2007) and SHH signaling, via its effector GLI1, has been implicated in the specification of this subtype (Ihrie et al., 2011). Interestingly, previous work demonstrated that Vax1 expression is positively controlled by SHH signaling (Hallonet et al., 1999; Take-uchi et al., 2003; Furimsky and Wallace, 2006). In light of our finding that Vax1 deletion also leads to specific loss of CB-N, it appears probable that Vax1 acts downstream of SHH expression in the ventral SVZ to control CB-N production for the OB.

In addition to this local role in CB-N generation, Vax1 regulates the generation of a neighboring neuron type. Indeed, its expression extends in a gradient far into dorsal regions of the ventricular wall, where Pax6 is expressed and acts as a key component of DA-N generation (de Chevigny et al., 2012a). Forced expression of Vax1 in the Pax6-positive domains was sufficient to reduce PAX6 protein expression and to inhibit the production of DA-N, but not of other neuron types, in the OB. This strongly indicates that Vax1 acts as a repressor of Pax6, comparable to the situation in the developing eye (Bertuzzi et al., 1999; Hallonet et al., 1999; Mui et al., 2005).

Loss-of-function of Vax1 through targeted electroporation with a CRE expression vector in the lateral wall of conditional mutants did not lead to a statistically significant increase in DA-N at the classically used confidence level of p ≤ 0.05. The observed tendency towards DA-N increase was, however, quite robust over several independent electroporation approaches implicating a large cohort of animals. We decided to include these data, as they complement the gain-of-function approach and as we believe that they could be biologically relevant. Indeed, the cell population in the lateral wall that will be able to induce DA-N fate after deletion of Vax1 is probably quite small. A large proportion of cells targeted by electroporation in the dorso-lateral wall do not express Vax1 and these cells will follow their normal differentiation program after recombination. Only cells in intermediate positions, that express sufficiently high Pax6 levels to be able to induce the DA-N phenotype while at the same time having sufficient Vax1 levels to suppress this differentiation pathway, will show a DA-N phenotype after Vax1 removal. In addition, other factors, like lower recombination efficiency of the Vax1 allele versus the R26tdTomato allele, might further diminish the number of cells in which the impact of gene inactivation can be studied (Long and Rossi, 2009; Luo et al., 2020). Thus, while our electroporation approach allows targeting and manipulating specific stem cell compartments, it also has limitations that restrict interpretation.

Finally, the implication of microRNAs introduces an additional level of complexity into the cross regulatory machinery that underlies OB interneuron fate determination. In previous work, we demonstrated that PAX6 protein, and consequently DA-N production, is confined to the dorsal aspect of the lateral ventricle wall by posttranscriptional regulation of the Pax6 3’UTR by the microRNA mir-7. The latter is, like Vax1, expressed in a ventro-dorsal and Pax6 opposing gradient. Thus, it is tempting to hypothesize that the repression of Pax6 by Vax1 is, at least in part, indirect and produced via activation of miR-7. Our finding that overexpression of Vax1 in vivo induced a strong increase in miR-7 levels strongly supports such a scenario.

## Material and Methods

### Animals

All animal procedures were carried out in accordance to the European Communities Council Directie 2010/63/EU and approved by French ethical committees (Comité d’Ethique pour l’expérimentation animale no. 14; permission numbers: 00967.03; 2017112111116881v2). Animals were held on a 12 h day/night cycle and had access to food and water ad libitum. Animals of both sexes were used for experiments. CD1 mice (Charles River, Lyon, France) were used for *in vivo* electroporation and expression pattern analyses. Vax1flox conditional mutants (Hoffman 2016) and Rosa26tdTomato reporter mice (Ai14, Jackson Laboratories, USA, RRID:IMSR_JAX:007914) were bred on a mixed C57BL/6*CD1 genetic background. *Vax1flox* genotyping was performed with *Vax1flox* forward: 5’-GCCGGAACCGAAGTTCCTA; *Vax1wt* forward: 5’-CCAGTAAGAGCCCCTTTGGG, reverse 5’-CGGATAGACCCCTTGGCATC. Ai14 genotyping was performed with *R26wt* forward: 5’-AAGGGAGCTGCAGTGGAGTA, reverse 5’-CCGAAAATCTGTGGGAAGTC; *R26tdTom* forward: 5’-CTGTTCCTGTACGGCATGG and reverse: 5’-GGCATTAAAGCAGCGTATCC.

### Plasmid and In vivo electroporation

The full-length rat cDNA sequence of Vax1 was excised from pCMV2-Rn-Vax1-FLAG (a gift of Kapil Bharti) and subcloned into pCAGGS vector to produce pCAG-Vax1. Post-natal day 0 (P0) or day 1 (P1) pups were electroporated as previously described (Boutin et al., 2008; de Chevigny et al., 2012a) with plasmid DNA or RNA (Bugeon et al., 2017). CRE Recombinase mRNA (130-101-113, a generous gift from A. Bosio, Miltenyi Biotec, Bergisch Gladbach, Germany) and pCX-CRE (Morin et al., 2007) were used at a concentration of 0.5 µg/µl. In CD1 pups, pCX-EGFP (Morin et al., 2007) was co-injected with pCAG-Vax1 or empty pCAGGS (as a control) in a 1:2 molecular ratio to label the electroporated cells. Targeting of the dorsal or lateral wall of the lateral ventricle was directed by distinct orientation of the electrodes. Brains were collected at different time points after electroporation.

### In situ hybridization and immunohistochemistry

For all procedures, tissues were fixed by intracardiac perfusion with ice-cold 4% paraformaldehyde (wt/vol) in phosphate buffered saline (PBS) and cryoprotected in 30% sucrose solution in PBS. Brains were sliced coronally using either a microtome (Microm Microtech, France) or a cryostat (Leica Biosystems, France).

Mouse Vax1 cDNA clone (GeneBank: BC111818, Imagene) was used to produce the anti-sense RNA probe. Combined *in situ* hybridization (ISH) and immunohistochemistry were performed as previously described (Tiveron et al., 1996) on 16 µm cryosections at post-natal day 3 (P3) using anti-digoxigenin antibody (Sheep polyclonal, 1:1000, Roche, 11093274910, RRID:AB_514497). Two days exposure to alkaline phosphatase (AP) substrate NBT/BCIP (Promega, S3771) was necessary to detect AP-DIG-labeled Vax1 mRNA at maximal level. Subsequently, antibodies directed against PAX6 (rabbit polyclonal, 1: 1000, Millipore, AB2237, RRID:AB_1587367), KI67 (mouse IgG1k clone B56, 1:200, BD Biosciences, 550609, RRID:AB_393778), ASCL1 (mouse, clone 24B72D11.1, 1:100, BD Biosciences, 556604, RRID:AB_396479), or DLX2 (rabbit polyclonal, 1: 2000 a gift from Prof. K. Yoshikawa) were applied to the ISH treated sections and were revealed with secondary antibodies coupled to horseradish peroxidase (Jackson ImmunoResearch Laboratories, UK), using 3, 3’-Diaminobenzidine (Invitrogen, 750118) as a substrate. Sections were finally mounted in fluoromount medium and analyzed by light microscopy using Axiovision Rel. 4.8 software (Zeiss, Germany).

For immunofluorescence, 50 µm floating sections were blocked in PBS supplemented with 0.5% Triton-X100, 10% foetal calf serum (FCS) and incubated overnight at 4°C in PBS, 0.1% Triton, 5% FCS with primary antibodies: anti-Calbindin D-28K (rabbit polyclonal, 1:1000, Millipore, AB1778, RRID:AB_2068336 or mouse IgG1, 1:3000, Swant, 300, RRID:AB_10000347), anti-Calretinin (mouse IgG1, clone 37C9, 1:1000, Synaptic systems, 214111, RRID:AB_2619906), anti-Tyrosine hydroxylase (chicken IgY, 1:1000, Avès labs, TYH, RRID:AB_10013440). After washing in PBS, Alexa Fluor-conjugated secondary antibodies (Jackson ImmunoResearch Laboratories) were applied diluted at 1:500 in blocking solution for 2 h at room temperature. After staining of cell nuclei with Hoechst 33258 (Invitrogen, H3569), sections were mounted with Mowiol (Sigma-Aldrich, 81381).

### RNA Isolation, qRT-PCR, cell dissociation, and FACS

Animals (P1-P3) were decapitated and brains were cut into 300 μm thick sections using a Vibrating-Blade Microtome (Thermo Scientific, HM 650V). V-SVZ and RMS tissues were micro-dissected under a binocular microscope and kept in cold Hank’s balanced salt solution (HBSS, Gibco, 14170120). RNA was extracted using the miRNAeasy kit (Qiagen, 217004) or by Trizol reagent (life technologies, 15596026) according to manufacturer instructions, allowing the recovery of long and short RNAs. For Pax6 and Vax1 expression analysis, cDNA was prepared using superscript III reverse transcriptase (ThermoFisher Scientific, 12574-030) following manufacturer instructions and quantitative PCR was performed on a BioRad CFX system using SYBR GreenER qPCR SuperMix (ThermoFisher Scientific, 11762100). ß-Actin was used as reference gene. Primers used for mRNA detection are the following: Beta Actin forward 5’-CTAAGGCCAACCGTGAAAAG and reverse 5’-ACCAGAG GCATACAGGGACA; Pax6 forward 5’-TGAAGCGGAAGCTGCAAAGAAA and reverse 5’-TTTGGCCCTTCGATTAGAAAACC; Vax1 forward 5’-GCTTCGGAAGATTGTAACAAAAG and reverse 5’-GGATAGACCCCTTGGCATC.

For miRNA expression, cDNA was prepared using miScript II (Qiagen, 218160) and qPCR was performed using miScript SYBR Green PCR Kit (Qiagen, 218073) together with miRCURY LNA miRNA PCR Assay (Qiagen, 339306) on a StepOne Real-Time PCR System (Applied Biosystems). Let-7a was used as normalizer miRNA.

To generate single-cell suspensions for FACS, dissected tissues were subjected to cell dissociation using a Papain solution and mechanical dissociation as described previously (Lugert et al 2010). Cells were resuspended in HBSS/Mg/Ca supplemented with 10mM HEPES (Gibco, 15630080), 40 μg/ml DNase I (Roche, 10104159001), 4.5 g/L Glucose (Gibco, A2494001) and 2mM EDTA, filtered through a 30 μm pre-separation filter (Miltenyi Biotec, 130-041-407) and sorted on MoFlo Astrios EQ cytometer (Beckman-Coulter) gating on GFP-positive population.

### Image analyses

All images were analyzed blind to the experimental condition. Optical images were acquired with an Axioplan2 ApoTome microscope (Zeiss, Germany) using ZEN software (Zeiss, RRID:SCR_013672), and processing was performed using Fiji software (RRID:SCR_002285, (Schindelin et al., 2012)) or ImageJ (NIH, https://imagej.nih. gov/ij/, RRID:SCR_003070).

### Statistical analysis

Data are presented as mean ± SD. Histograms were drawn with GraphPad Prism version 8 (GraphPad Software, San Diego, California USA). Statistical analyses were performed using R software (RRID:SCR_001905) and R Commander Package (https://CRAN.R-project.org/package=Rcmdr). The non-parametric two-tailed Mann Whitney U test was performed for all in vivo experiments. Differences were considered statistically significant when p ≤ 0.05.

## Acknowledgement

We thank the members of the Cremer lab for support and critical reading of the manuscript. We thank the local PiCSL-FBI core facility (IBDM, AMU-Marseille) supported by the French National Research Agency through the « Investments for the Future’ program (France-BioImaging, ANR-10-INBS-04) as well as the IBDM animal facility. We are grateful to AMUTICYT Cytometry and Cell Sorting Core facility, AMU, UMR-S 1076. This work was supported by the Agence National pour la Recherche (grants ANR-13-BSV4-0013 and ANR-17-CE16-0025), Fondation pour la Recherche Medicale (FRM) “Label Equipe FRM” and Fondation de France (FDF) grant FDF70959 to HC. AE was supported for a postdoctoral fellowship from the Swiss National Funds. NIH grants to PLM in support of this work are P50 HD12303, R01 HD072754, R01 HD082567, P30 CA23100, P30 DK063491, and P42 ES010337. HMH was supported by NIH K99 HD084759.

---

**Figure 4 - figure supplement 1.**
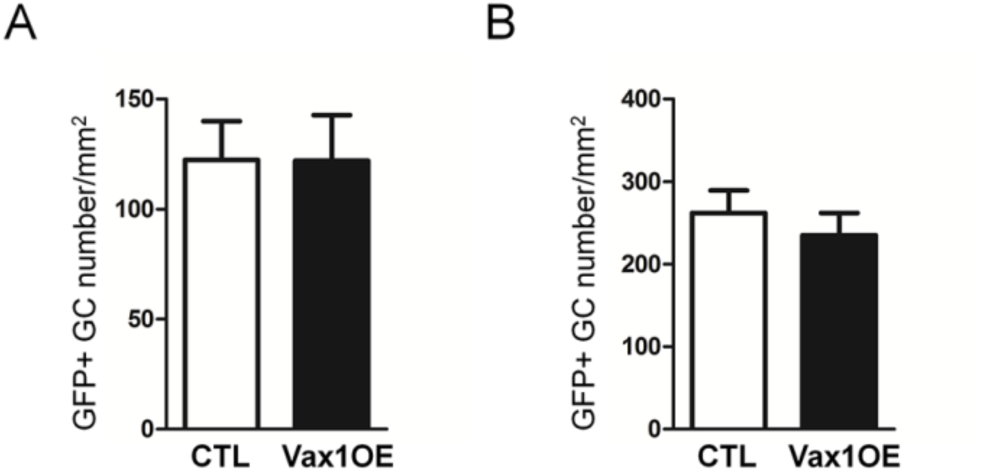
Forced expression of Vax1 has no effect on cell density in the OB granule cell (GC) layer, 15 days after electroporation of dorsal (CTL n=15, Vax1 n=14) or (B) lateral (n=10 for both conditions) V-SVZ progenitors. Data are represented by mean ± SEM.

